# Polygenic scores via penalized regression on summary statistics

**DOI:** 10.1101/058214

**Authors:** Timothy Shin Heng Mak, Robert Milan Porsch, Shing Wan Choi, Xueya Zhou, Pak Chung Sham

## Abstract

Polygenic scores (PGS) summarize the genetic contribution of a person’s genotype to a disease or phenotype. They can be used to group participants into different risk categories for diseases, and are also used as covariates in epidemiological analyses. A number of possible ways of calculating polygenic scores have been proposed, and recently there is much interest in methods that incorporate information available in published summary statistics. As there is no inherent information on linkage disequilibrium (LD) in summary statistics, a pertinent question is how we can make use of LD information available elsewhere to supplement such analyses. To answer this question we propose a method for constructing PGS using summary statistics and a reference panel in a penalized regression framework, which we call lassosum. We also propose a general method for choosing the value of the tuning parameter in the absence of validation data. In our simulations, we showed that pseudovalidation often resulted in prediction accuracy that is comparable to using a dataset with validation phenotype and was clearly superior to the conservative option of setting the tuning parameter of lassosum to its lowest value. We also showed that lassosum achieved better prediction accuracy than simple clumping and *p*-value thresholding in almost all scenarios. It was also substantially faster and more accurate than the recently proposed LDpred.

## Introduction

A vast number of twin studies as well as recent genome-wide association studies have demonstrated that a large proportion of the variance in liability to common diseases and human traits is due to genetic differences between individuals (Polderman *et al*., 2015; Yang *et al*., 2011; Bulik-Sullivan *et al*., 2015). These studies have also made clear that only a very small proportion of the total genetic contribution can be unambiguously attributed to variation in particular loci of the genome. The vast majority of such genetic contribution is thus spread across the huge landscape of the genome, with many loci each contributing a small, almost undetectable effect on the phenotypes (Dudbridge, 2013, 2016). One important source of evidence towards this conclusion is from studies that examined the association of polygenic predictors of diseases/traits, where it has been repeatedly found that SNPs that are not themselves significantly associated with the phenotypes can, by being aggregated as a score, be very significantly associated with the phenotypes in different samples (Agerbo *et al*., 2015; Byrne *et al*., 2014; Evans *et al*., 2009; Wei *et al*., 2009; Purcell *et al*., 2009; Ripke *et al*., 2013; Speliotes *et al*., 2010; Machiela *et al*., 2011; Stahl *et al*., 2012; Martin *et al*., 2015; Chang *et al*., 2014). A particular remarkable demonstration is that persons with such *polygenic scores* for schizophrenia at the top 10 percentile of the population can be at more than 10 times the risk of having the disease than those at the bottom 10 percentile (Ripke *et al*., 2014; Agerbo *et al*., 2015), raising hope that one day a person’s risk for many common disease can be accurately assessed simply by the examination of one’s genome.

Thus, there is considerable interest in the calculation of such polygenic scores (PGS) in GWAS and Genome-wide meta-analyses, where they are also known as risk scores (Ripke *et al*., 2013; Domingue *et al*., 2014), polygenic risk scores (e.g. Euesden *et al*., 2015; Byrne *et al*., 2014; Agerbo *et al*., 2015; Dudbridge, 2013), and allelic scores (Burgess and Thompson, 2013; Evans *et al*., 2013). In a typical application, a unique PGS is assigned to each individual based on the person’s genotype. The score summarizes the genetic contribution to a particular disease or phenotype for that indivdiual given his/her genotype. They are then used for testing of complex genetic contribution due to multiple loci or even the entire genome, or the examination of genetic correlation, or are used as a covariate for the adjustment of genetic effects in a multiple regression model (Wray *et al*., 2014).

From a statistical perspective, polygenic scores are weighted sums of the genotypes of a set of SNPs. In most applications of PGS, the weights are usually the SNPs’ individual regression coefficients with the phenotype (e.g. Purcell *et al*., 2009; Wray *et al*., 2014; Euesden *et al*., 2015). A critical issue is the total number of SNPs that should be included in the PGS. Although it is usually advisable to use a liberal *p*-value cutoff in the selection of SNPs to be included, the optimal *p*-value cutoff is generally unknown (Wray *et al*., 2014). As a result, in many studies, PGS are constructed using a number of thresholds (Purcell *et al*., 2009; Ripke *et al*., 2014; Byrne *et al*., 2014; Martin *et al*., 2015; Chang *et al*., 2014), and there is at least one piece of software developed to facilitate this (Euesden *et al*., 2015). Generally, we focus on the *p*-value threshold that achieves the highest correlation/association with the phenotypes in a validation dataset that contains a measure of the phenotype under study. This approach, however, becomes less useful if the phenotype is not available in the target dataset. Recently, Mak *et al*. (2016) sought to overcome this problem by downweighting the usual weights by the SNPs’ local true discovery rate, where the additional downweighting or shrinkage factor can be estimated using a data-driven approach. Although *p*-value thresholds were not needed, they showed that this leads to comparable predictive performance with the best *p*-value threshold.

Another issue with this standard approach to PGS calculation is that there is no account taken of the fact that SNPs are in linkage disequilibrium (LD) with each other. If SNPs of a particular locus which are in high LD with one another are all included in the score, the contribution to the PGS due to that locus will be exaggerated in the score. For this reason, it is often recommended that SNPs be *pruned* before the application of PG scoring, such that highly correlated SNPs within a locus will have one or more removed (Purcell *et al*., 2009). Such an approach, however, may well reduce the predictive power of the PGS, as SNPs that are most predictive of the phenotype may be pruned away. A recent suggestion which has become very popular is that of clumping, which selectively removes less significantly related SNPs to reduce LD (Wray *et al*., 2014).

In principle, various machine learning methods or Bayesian methods can be applied in the construction of PGS, as they have been applied in the estimation of breeding values in animal studies (Meuwissen *et al*., 2001; Abraham *et al*., 2013; Szymczak *et al*., 2009; Habier *et al*., 2011; Pirinen *et al*., 2013; Erbe *et al*., 2012; Ogutu *et al*., 2012; Zhou *et al*., 2013). These methods do not require the assumption of SNP independence or near independence, and have been shown to perform better than simple PGS in simulation settings. However, their disadvantage is that they cannot be applied to summary statistics. Researchers without access to large datasets are thus unable to take advantage of the power offered by these studies or meta-analyses. A recent development in this direction is Vilhjálmsson *et al*. (2015). The authors proposed an approximate Bayesian method known as LDpred that calculates PGS based on summary statistics, using LD information from a reference panel. Such a development is particularly welcome due to the ready availability of summary statistics from many consortia, often calculated from tens to hundreds of thousands of individuals.

In this paper, we present an alternative method based on penalized regression. It is a deterministic method and a convex optimization problem, and as such does not suffer from problems of non-convergence, which is a possible problem with LDpred. It is also substantially faster than LDpred, and in our simulations achieved near-best prediction performance across a wide variety of scenarios. As a side observation, it was also found that LDpred did not achieve the improved prediction performance claimed by the authors in our simulations. As with any machine learning approach, our proposed method requires the choice of a tuning parameter. This is particularly difficult when we do not have raw data and hence cannot perform cross-validation. Here, we offer a solution that can potentially be applied more generally. The approach is presented in the methods section and we assessed its performance by simulation studies. Insights gained from the simulations are discussed.

## Material and methods

### The LASSO problem in terms of summary statistics

Given a linear regression problem

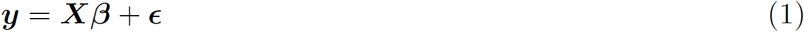
where ***X*** denotes an *n*-by-*p* data matrix, and ***y*** a vector of observed outcomes, the LASSO (Tibshirani, 1996) is a popular method for deriving estimates of *β* and predictors of (future observations of) ***y***, especially in the case where *p* (the number of predictors/columns in ***X***) is large and when it is reasonable to assume that many *β* are zero. LASSO obtains estimates of *β* (weights in the linear combination of ***X***) given ***y*** and ***X*** by minimizing the objective function

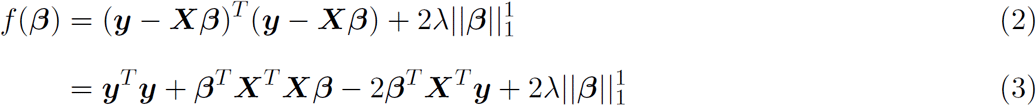
where 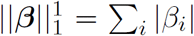 denote the *L*_1_ norm of *β*, for a particular fixed value of *λ*. In general, depending on *λ*, a proportion of the *β_i_* are given the estimate of 0. It is also a specific instance of *penalized regression* where the usual least square formulation of the linear regression problem is augmented by a penalty, in this case 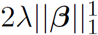. LASSO lends itself to being used for estimation of ***β*** in the event where only summary statistics are available, because if ***X*** represent standardized genotype data and ***y*** standardized phenotype, divided by 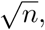, then equation (3) can be written as:

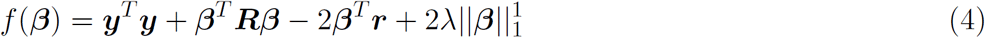
where ***r*** = ***X***^*T*^***y*** represents the SNP-wise correlation between the SNPs and the phenotype, and ***R*** = ***X***^*T*^ ***X*** is the LD matrix, a matrix of correlations between SNPs. As we can obtain estimates of ***r*** from summary statistics databases that are publicly available for major diseases/phenotypes (see e.g. the list from Pasaniuc and Price, 2016) and LD hub (http://ldsc.broadinstitute.org/), and estimates of LD (***R***) from publicly available genotype such as the 1000 Genome database (1000 Genomes Project Consortium, 2015), equation (4) suggests a method for deriving PGS weights as estimates of ***β*** by minimizing *f*(***β***).

An issue that surfaces when we substitute ***R*** and ***r*** with the estimates derived from publicly available data is that the genotype ***X*** used to estimate ***R*** and ***r*** will in general be different. In particular, it will be more appropriate to write 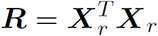 to indicate that the genotype used to derive estimates of LD (***X***_*r*_) will not in general be the same as the genotype that gave rise to the correlations ***r***. Writing equation (4) as

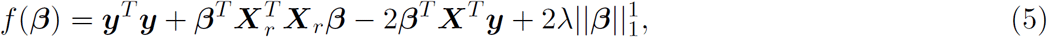
however, would imply that (5) is no longer a LASSO problem, because it is no longer a penalized least squares problem. A minimum to (5) can still be sought, although the solutions would often be unstable and non-unique, since 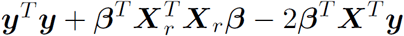 will not generally have a finite minimum.

A natural solution to this problem is to *regularize* equation (5). In particular, if we replace 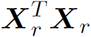 with 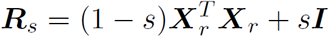, for some 0 < *s* < 1, then

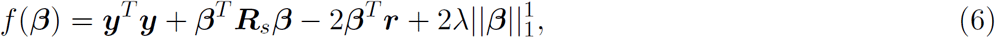
will be equivalent to a LASSO problem.

*Proof*. First, we note that ***y***^*T*^***y*** is a constant and thus replacing it with any other constant will not change the solution. ***R***_*s*_ is necessarily positive definite for 0 < *s* < 1. This means that there always exists ***W*** and ***v*** such that

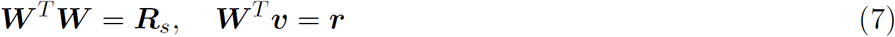

Substituting (7) into (6) and replacing ***y***^*T*^***y*** with ***v***^*T*^***v***, we see that (6) can be written in a form such as (2) and is therefore a LASSO problem.

Expanding (6) into

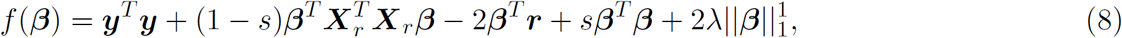
we note that (8) encompasses a number of submodels as special cases. For example, when *s* = 1, estimates of ***β*** will be equivalent to applying a “soft” threshold to the univariate correlation summary statistics ***r*** (as opposed to the “hard” thresholds using *p*-values.) In particular,

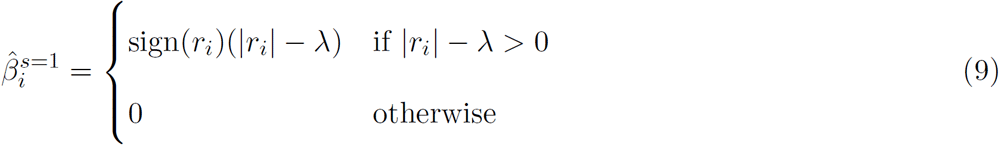

(Zou and Hastie, 2005). Note that because there is a monotonic relationship between univariate *p*-values and unsigned correlation coefficients (coming from the monotonic relationship between correlation coefficients and *t*-statistics with *n* − 2 degrees of freedom, equation (15)), soft-thresholding using correlation coefficients can be expected to be very similar to *p*-value thresholding. Another feature is that when *λ* = 0, the problem is similar to applying ridge regression to estimate ***β***, except for a constant scaling value. In most cases, the scale of a PGS is irrelevant, since it is almost never directly used in genomic risk prediction without appropriate scaling (e.g., in So *et al*., 2011). For a particular choice of *s*, therefore, equation (8) results in genomic BLUP (Best Linear Unbiased Predictors) (de Los Campos *et al*., 2013). When *λ* = 0 and *s* = 1, the estimated PGS becomes equivalent to simply using the entire set of correlation estimates without shrinkage or subset selection.

Moreover, (8) is simply an elastic net problem (Zou and Hastie, 2005), and thus can be solved using fast coordinate descent algorithms (Friedman *et al*., 2010) for many values of *λ* at a time. In particular, using this algorithm, it is not necessary to compute the *p*-by-*p* matrix 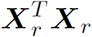, which would be extremely memory-consuming even for tens of thousands of SNPs. Denoting 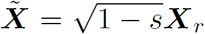 equation (8) can be obtaining by iteratively updating *β_i_* as

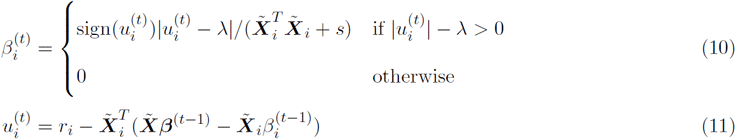

A more detailed proof of equations (10) and (11) is given in the Supplementary materials. An R package that carries out the estimation of ***β*** is made available at https://github.com/tshmak/lassosum. We made special effort to allow estimation to be done directly on PLINK 1.9 (Chang *et al*., 2015) .bed files, eliminating the need to load large genotype matrices into R.

### Selection of tuning parameters

As with standard elastic net problems, in any application, *λ* and *s* need to be chosen. Generally, in the presence of a validation dataset, we can choose *λ* by maximizing the correlation of the PGS with the validation phenotype data, just as it has been done in the choice of a *p*-value cutoff points in standard PGS calculations (Wray *et al*., 2014; Euesden *et al*., 2015). In principle we can use this method to choose a suitable value for *s* also, although repeating the estimation over different values of *s* is much more time-consuming. Thus in this paper we set *s* to a few chosen values and examined whether they are sufficient in arriving at a PGS with reasonable prediction accuracy.

A more pressing problem is that validation phenotypes are not often available. And here we try to simulate this procedure in the following manner, which we refer to as *pseudovalidation* in this paper. First, note that the correlation between a 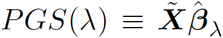 and the phenotype 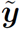 in a new “test” dataset with standardized genotype 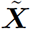

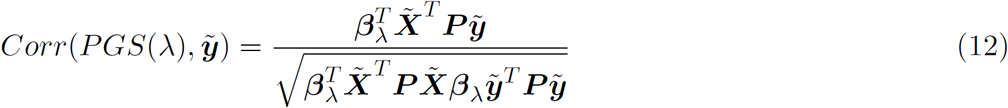
where ***P*** = ***I*** − **11**^*T*^/*n* is the mean-centering matrix.

In the absence of validation data, 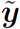 is unavailable. Our solution is to substitute 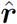 for 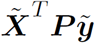, where 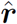 is a shrunken estimate of the ***r***, the observed correlation coefficient vector. Since 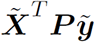 can be interpreted as a correlation coefficient only if 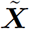 is a standardized genotype matrix and 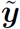 standardized phenotype, we replace 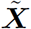 with its standardized version, 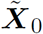, and discard the constant 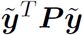 term, so as to maximize the function

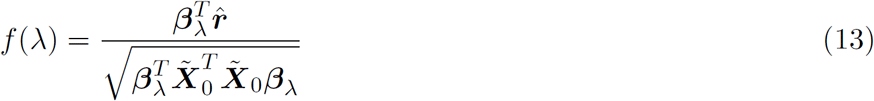
over *λ*. Here, following Mak *et al*. (2016), we calculated

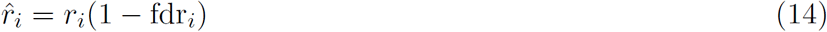
where fdr_*i*_ is the local false discovery rate of SNP *i*. While Mak *et al*. (2016) estimated fdr*i* using maximum likelihood and a non-parametric kernel density estimator, we found that Strimmer (2008) provided a fast, nonparametric estimator for fdr*i* which is constrained to be monotonic decreasing with |*r*_*i*_|, and it is this approach that we have implmented in the simulations.

### Some notes on application

In the above, we have assumed that the SNP-wise correlations (***r***) will be available from the summary statistics. When these are not available, we suggest *pseudo-correlation* estimates 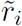 be derived by converting *p*-values to correlation, using the monotonic relationship between *t*-statistics and correlations:

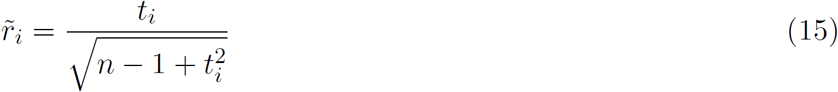

In our simulations this resulted in almost identical estimates as using actual (Pearson’s product moment) correlations (Figure S1).

Another issue is that in the theory given above, we assume that ***X*** and ***y*** have been standardized such that ***r*** represent the correlation coefficients between the genotype and the phenotype. We note that such standardization can be justified by the fact that the LASSO is often performed on standardized variables (Li *et al*., 2012; Hastie *et al*., 2009; Yi *et al*., 2014). However, when it comes to the construction of PGS, we ought to use unstandardized coefficients as weights. To convert standardized coefficients to unstandardized ones, we can simply use the formula

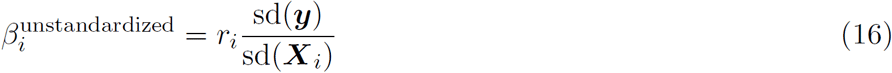

However, since sd(***y***) and sd(***X***_*i*_) are generally unavailable, we can use 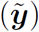 and 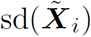 from the validation data instead. Using these also prevents any SNP from undue influence in the overall PGS due to the division of 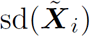 close to 0, since a SNP’s variance contribution is proportional to its variance and the square of the coefficients.

A third issue concerns the difference between the SNPs with summary statistics and the SNPs that are included in the reference panel. Often the reference panel may not contain all SNPs with summary statistics. Equivalently, there may be no variation within the panel for some SNPs. In LDpred, these SNPs are discarded by default. However, we think that this is not necessary, as it may result in the removal of SNPs that are predictive of the disease/phenotype. An intuitive approach to dealing with these SNPs is that we treat them as if they were all mutually independent and apply soft-thresholding as in (9). Equivalently, we let ***X***_*ri*_ causal SNPs across the genome for these SNPs to be a vector of zero, and we augment equation (8) by a term 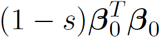

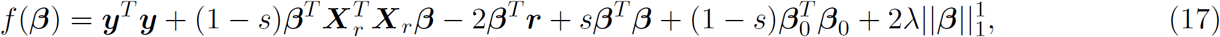
where ***β***_0_ denotes the sub-vector of ***β*** whose sd(***X***_*i*_) = 0, such that the total ridge penalty for these parameters is 1.

A fourth issue concerns the application of pseudovalidation to clumped data. We proposed above that 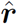 be estimated using (14) and that the local false discovery rates be estimated using the procedure of Strimmer (2008). An important point is that the method assumes that a sizeable proportion of the ***r*** are in fact null. Under clumping, this may not necessarily be the case, and we therefore suggest estimating fdr_*i*_ and hence 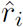 *before* applying clumping.

### Simulation studies

We performed a number of simulation studies to assess the performance of our proposed method, which we refer to in this paper as 

~~~
lassosum
~~~

. In our first simulation study, we made use of the Welcome Trust Case Control Consortium (WTCCC) Phase 1 data for seven diseases. We filtered variants and participants using the following QC criteria: genotype rate > 0.99, Minor Allele Frequency > 0.01, Missing genotype per individual < 0.01, SNP rsID included in the 1000 Genome project (Phase 3, release May 2013) genotype data, with matching reference and alternative alleles, on top of the QC done by the original researchers (Wellcome Trust Case Control Consortium, 2007). This resulted in 358,179 SNPs and 15,603 individuals, of which 2,859 were controls. In our first set of simulations, we ignored the phenotype data and generated our own based on the linear model

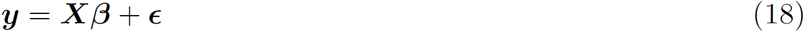
where ***X*** is the unstandardized genotype matrix, and *ϵ* ∼ *N*(0, *σ*^2^***I***) represents random error. The distribution of the causal effects ***β*** ≡ *vec*({*β_i_*}) ≡ *vec*({*β*_*jk*_}) is generated using a similar scheme to that described in Vilhjálmsson *et al*. (2015):

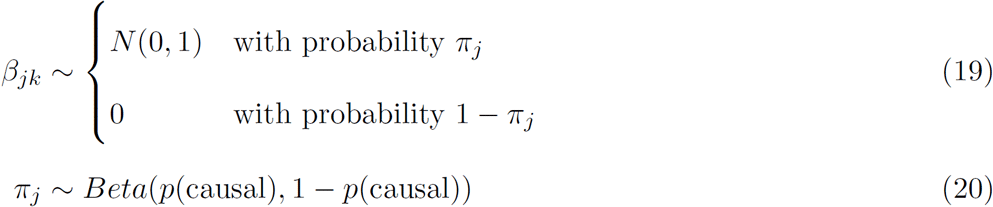
where *j* denotes genomic regions and *k* indices SNPs within the region and *i* is a general index for all SNPs in the database, and *p* is the expected proportion of causal SNPs across the genome (note *E*(*π_j_*) = *p*(causal)). Genomic regions were defined using the 1,725 LD blocks obtained from the 1000 Genomes European (EUR) sub-population, as provided by Berisa and Pickrell (2015).

We derived standardized ***β*** as

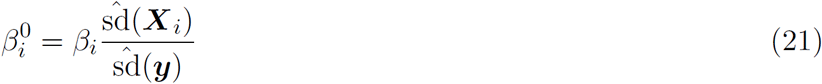
and observed correlation coefficients as

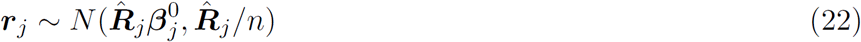
where 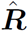 is the observed correlation matrix of the *j*th region from the genotype ***X*** and *n* is the sample size. We set 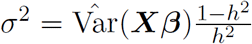 and *h*^2^ = 0.5 in our calculation of ***y***.

We randomly chose two 1,000 samples as two test datasets. In the first dataset ***X***^(1)^, validation and pseudovalidation were performed to determine the optimal value of *λ*. This choice of *λ* and/or *s* was applied in the other test dataset ***X***^(2)^ in the assessment of prediction accuracy. Prediction accuracy was assessed by the correlation of the PGS with the true predictor ***X***^(2)^***β***. Except when assessing the performance of using different reference panels, we used the first test dataset ***X***^(1)^ as the reference panel also.

In assessing of the impact of using different reference panels, we let the 1000 Genome East Asian (EAS) sub-population (*n* = 503) be our test dataset. We compared the performance of using four different reference panels: (1) the original sample that generated the summary statistics, (2) a sample of 1,000 from the WTCCC, (3) the European sub-population from the 1000 Genome project, and (4) the East-Asian sub-population from the 1000 Genome project.

The above simulations were repeated 10 times and were compared with the approach of *p*-value thresholding (with and without clumping) and LDpred. For clumping, we used a window of 250 kb and an *R*^2^ of {0.1, 0.2, 0.5, 0.8}. (See Supplementary Note for a brief explanation of clumping.) For *p*-value thresholding, we used the set of *p* values {5*e*^−8^, 1*e*^−5^, 1*e*^−4^, 1*e*^−3^, 0.0015, 0.002, 0.0025, …, 0.995, 1} as possible *p*-value thresholds. For LDpred, we used the set of proportion of causal SNPs {0.001, 0.003, 0.01, 0.03, 0.1, 0.3, 1}. The size of the window for LD calculation was calculated as the number of SNPs in the dataset divided by 3,000, as recommended in the LDpred paper. For *p*-value thresholding and LDpred, we used a validation dataset as well as pseudovalidation to select the best threshold and proportion of causal SNPs, respectively.

Because summary statistics are often calculated from large sample sizes and for a large number (often around 10 million) of SNPs, we also attempted to carry out simulations using a larger dataset. In particular, we wanted to see whether clumping is an efficient strategy for data reduction, as the speed of 

~~~
lassosum
~~~

 suffers with such a large number of SNPs. For this purpose, we first identified SNPs from the summary statistics derived in the meta-analysis of Okada *et al*. (2014) for Rheumatoid Arthritis (RA) that were common with those in the 1000 Genome dataset. We then generated our own summary statistics using the above method (equations 18 to 22), using the European (EUR) subsample of the 1000 Genome dataset as a base. This resulted in a dataset of 8,270,298 SNPs. We used the EUR subsample as the reference panel and the East Asian (EAS) subsample of the 1000 Genome dataset as the test sample to assess the predictive performance.

Finally, we assessed the performance of 

~~~
lassosum
~~~

 using real summary statistics from large meta-analyses. Summary statistics were downloaded from five publicly available resources: Bipolar disorder (https://www.med.unc.edu/. Sklar *et al*. (2011), *n*(cases) = 7, 481, *n*(controls) = 9, 250), Coronary Artery Disease (http://www.cardiogramplusc4d.o. Nikpay *et al*. (2015), *n*(cases) = 60, 801, *n*(controls) = 123, 504), Crohn’s disease (http://ibdgenetics.org/downloads. Liu *et al*. (2015), *n*(cases) = 22, 575, *n*(controls) = 46, 693), Rheumatoid Arthritis (http://plaza.umin.ac.jp/˜yokada Okada *et al*. (2014), *n*(cases) = 14, 361, *n*(controls) = 43, 923), and Type 2 diabetes (http://diagram-consortium.org/, Mahajan *et al*. (2014), *n*(cases) = 26, 488, *n*(controls) = 83, 964). The performance of PGS derived using 

~~~
lassosum
~~~

 and other methods were assessed using the WTCCC data. Because all of these meta-analyses included the WTCCC as one of the studies, PGS derived using these summary statistics directly would overfit the data. To overcome this problem, we attempted to isolate the non-WTCCC components of the summary statistics by reversing the fixed-effects meta-analysis equations:

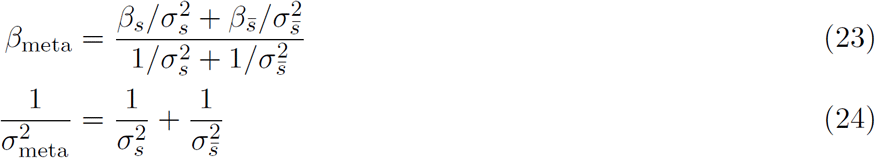
where *β_s_* and *σ_s_* denote the log odds ratio and standard error from the WTCCC study and 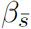 and 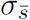 the contribution to the meta-analysis apart from WTCCC. SNPs with negative 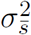 were set to have zero effect size. *p*-values were derived from 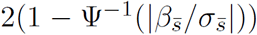 and converted to correlations using (15). Prediction accuracy of the summary statistics-derived PGS were assessed by the Area Under the ROC Curve (AUC) statistic when used to predict disease status in the WTCCC dataset with the relevant disease and the 2,859 controls. The testing sample was also used as the reference panel.

In all of the above analyses, we carried out estimation by LD blocks as defined by Berisa and Pickrell (2015).

## Results

Our WTCCC simulations were performed with summary statistics sample sizes of 10,000, 50,000 and 250,000 respectively. We used two values for *p*(causal), the expected proportion of causal SNPs: 0.1 and 0.01. *p*(causal) = 0.01 represents a scenario where there are fewer causal SNPs and effect sizes are larger. Conversely *p*(causal) = 0.1 represents a scenario where causal SNPs have smaller effect sizes and are more spread out over the genome. Figure S2 displays the performance of 

~~~
lassosum
~~~

 with different values of *λ* for one of the simulations. It can be seen that in all the simulation scenarios, the general pattern is that predictive performance increases with *λ* up to a point and then decreases, often rapidly. Using a validation dataset or alternatively pseudovalidation is usually effective in helping us select a value of *λ* that is close to the optimal. Comparing different values of *s*, the shrinkage parameter, we see that the maximum attainable correlation is generally lower for *s* = 1, the scenario where 

~~~
lassosum
~~~

 reduces to soft-thresholding, i.e. where information on LD is ignored, except when *n* = 10000 and *p*(causal) = 0.1. In addition, *s* = 0.5 and *s* = 0.2 usually gives better performance than *s* = 0.9. In Figure 1A, we give the average prediction performance over 10 simulations, comparing the use of pseudovalidation and a validation dataset with phenotype data as well as using the minimum *λ* value of 0.001. We use *λ* = 0.001 for comparison because it is shown in Figure S2 that in general the prediction performance of 

~~~
lassosum
~~~

 approaches a constant as *λ* tends to 0, whereas when *λ* approaches 1, the performance drops sharply. Thus, using *λ* close to 0 represents a conservative, safe option, and as noted before *λ* = 0 is equivalent to ridge regression. When *s* = 0.2 or 0.5, the performance of pseudovalidation was very similar to using a real validation phenotype. Both approaches were clearly superior to the conservative option of setting *λ* = 0.001. When *s* = 0.9 or *s* = 1, pseudovalidation was still clearly superior to setting *λ* = 0.001 for *n* = 10000 and *n* = 50000 and *p*(causal) = 0.01. In all simulations, the performance of *p*-value thresholding was similar to the use of 

~~~
lassosum
~~~

 with *s* = 1. Thus “soft-thresholding” and “hard-thresholding” appeared to give similar performance. We also observed that 

~~~
lassosum
~~~

 with *s* = 0.2 or *s* = 0.5 tended to give the best performance overall. In our implementation of 

~~~
lassosum
~~~

, the computation time for *s* = 0.2, 0.5 and 0.9 were similar (Figures S4 and S5). Thus, it is reasonable to maximize over *s* also using either a validation phenotype or pseudovalidation when using 

~~~
lassosum
~~~

. In Figure 1B, we compare the performance of 

~~~
lassosum
~~~

 with clumping and *p*-value thresholding, as well as with LDpred. For 

~~~
lassosum
~~~

, we optimized over both *λ* and *s* = {0.2, 0.5, 0.9, 1}. For comparison, we optimized over *p*-value thresholds and clumping *R*^2^ = {0.1, 0.2, 0.5, 0.8, No clumping}. For LDpred, we optimized over *p*(causal) = {0.001, 0.003, 0.01, 0.03, 0.1, 0.3, 1}. For *p*-value thresholding, clumping led to a noticeable increase in prediction accuracy, except when *p*(causal) = 0.1 and *n* = 10000. However, in all scenarios, 

~~~
lassosum
~~~

 was superior to clumping and thresholding. The result was similar whether the method was optimized using a validation dataset or pseudovalidation. We found that LDpred did not appear to have the claimed advantage over *p*-value thresholding in our simulations. At first we thought this might be because the size of the reference sample used was only 1,000, smaller than the recommended size of at least 2,000 in the paper. However, we found that the performance of LDpred did not improve even when the sample size of the reference panel (and test panels) were set to 5,000 (Figure S3).

A possible criticism of our simulations so far is that we performed 

~~~
lassosum
~~~

 by LD blocks defined by Berisa and Pickrell (2015), while the summary statistics were also simulated by the same LD blocks. To address this issue, we repeated the analysis using blocks with roughly the same number of SNPs spread uniformly across the genome. The number of blocks were made equal to the number of blocks given by Berisa and Pickrell (2015), but the boundaries were different. This would allow 

~~~
lassosum
~~~

 to adjust for LD within blocks but not LD across blocks in the boundary regions. We also compared it to the scenario when 

~~~
lassosum
~~~

 was carried out by chromosomes. The results are presented in Figure S4. It can be seen that 

~~~
lassosum
~~~

 by LD blocks and uniform blocks had nearly identical predictive performance. Thus the advantage that 

~~~
lassosum
~~~

 had in our simulations by sharing the same blocks by which the summary statistics were generated was negligible. The relative poor performance of 

~~~
lassosum
~~~

 when carried out by chromosomes is likely due to confounding by chance correlations between SNPs over long distances that are not in fact in LD.

In Figure 1C, we examined the effect of using different reference panels when using 

~~~
lassosum
~~~

. We generated the summary statistics using the entire WTCCC sample, and used four different reference panels for our LD information: (1) the original WTCCC sample that generated the summary statistics, (2) a sample of 1,000 from the WTCCC, (3) the European sub-population from the 1000 Genome project, and (4) the East-Asian subpopulation from the 1000 Genome project, which also served as the test sample. It was found that for the small sample size (*n* = 10, 000) scenario the use of the different reference panels made relatively little difference to predictive performance. However, as sample size increased, using the true sample that generated the summary statistics led to noticeably improved predictive performance. For many scenarios, using the 1000 Genome EUR sample as the reference panel led to a similar performance as using the original summary statistic sample. A clear advantage for using the summary statistics sample was only shown in the scenario with the most power (*n* = 250000 and *p*(causal) = 0.01). Using the wrong (EAS) reference sample was clearly inferior when the sample size was above 50,000, but it was still better than simple *p*-value thresholding.

Next we examined the performance of 

~~~
lassosum
~~~

 in a larger simulated dataset with around 8 million SNPs, with a focus on clumping, to see whether pre-filtering by clumping can be an effective method in reducing the number of SNPs in the analysis. The sample size for the summary statistics was set to 200,000. Six levels of clumping (*r*^2^ = 0.01, 0.05, 0.1, 0.2, 0.5, and 0.8) were applied to the data, using a window size of 250kb, resulting in around 190,000, 330,000, 430,000, 610,000, 1,170,000, and 1,940,000 SNPs respectively. (The actual number depends on the simulations.) We did not perform LDpred for *r*^2^ > 0.2 because it was too time and memory intensive. In Figure 2A, we present the results from this simulation. Here, we see that clumping was beneficial in improving prediction performance for *p*-value thresholding, and the best performance was achieved with an *r*^2^ of 0.5 or 0.8. For 

~~~
lassosum
~~~

, performance decreased with increasing level of clumping (decreasing *r*^2^). 

~~~
lassosum
~~~

 with no clumping gave the best performance overall. LDpred performed poorly in this simulation, likely because the reference panel size was too small.

**Figure 1:**
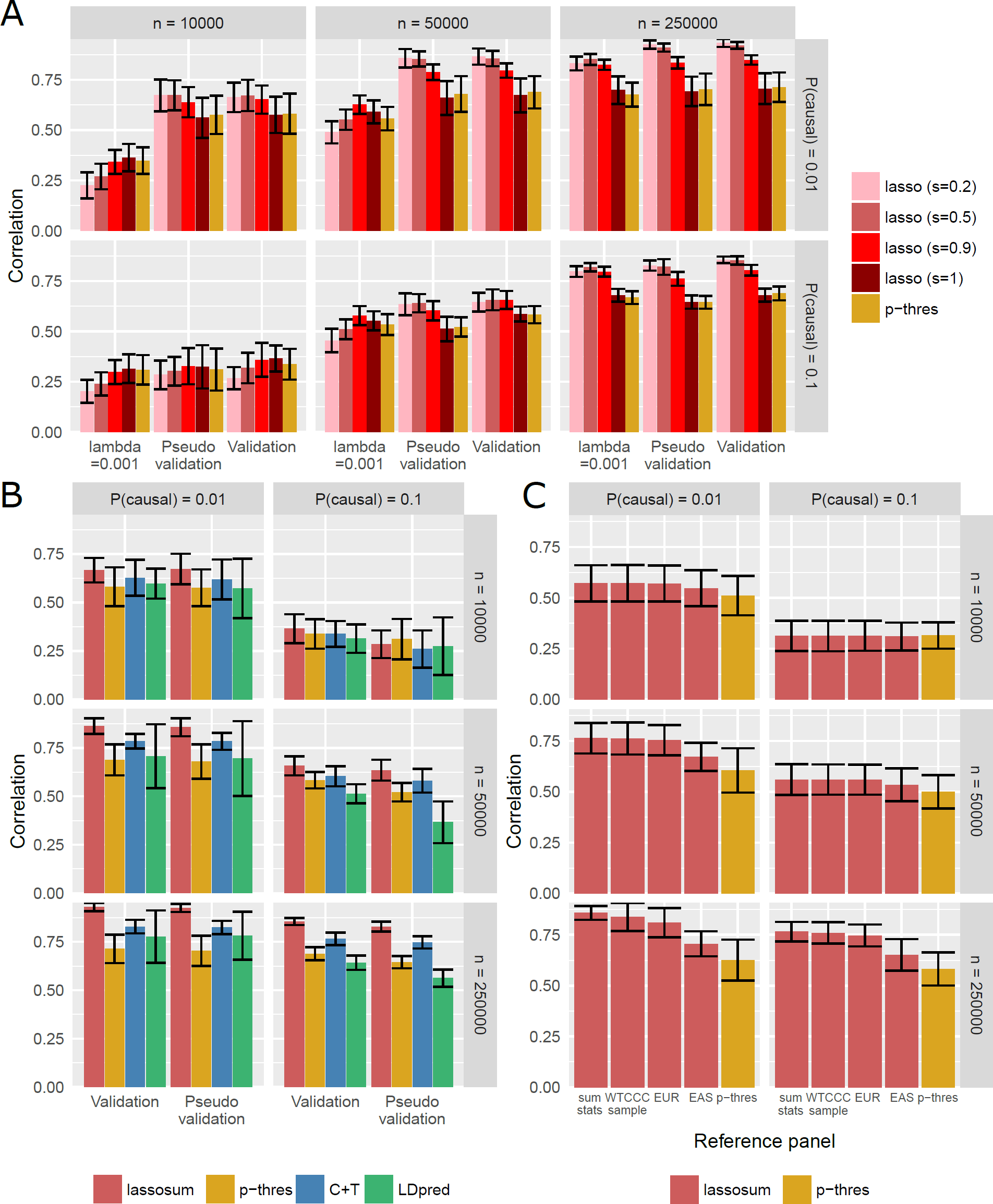
In all of the plots, mean and standard deviation of the correlation of the PGS with the true predictor are plotted. (A) Comparing the use of a validation dataset with phenotype data and pseudovalidation in selecting the tuning parameter *λ*. (B) Comparing the performance 

~~~
lassosum
~~~

, *p*-value thresholding (p-thres), *p*-value thresholding with clumping (C+T), and LDpred. (C) The effect of using different reference panels on 

~~~
lassosum
~~~

. sum stats: The same data from which the summary statistics were simulated, WTCCC sample: A sample of 1,000 from the WTCCC, EUR: European, EAS: East Asian reference panel from 1000G.

In Figure 2B, we present the results for using real summary statistics from five large meta-analyses to predict phenotypes in the WTCCC data. In all cases, the use of pseudovaliation resulted in a PGS that is close to the maximum AUC across all tuning parameters, and was clearly superior to using *λ* = 0.001. For BD, CAD, CD, and RA, the performance of 

~~~
lassosum
~~~

, LDpred, and clumping and thresholding were similar, although a slightly higher AUC was observed for 

~~~
lassosum
~~~

. For T2D, the maximum AUC was surprisingly achieved by *p*-value thresholding without clumping.

In Figures S5 and S6, we plot the average time taken to run 

~~~
lassosum
~~~

 on our computer cluster, using 1 core for each analysis. In general, running times for different values of *s* were similar, although lower values of *s* led to slightly longer running times. However, running times increased exponentially both with the number of participants (Figure S5) and the number of SNPs (Figure S6). Nonetheless, it was still substantially faster than LDpred. While LDpred typically requires hours to run, 

~~~
lassosum
~~~

 took only minutes.

## Discussion

In this paper, we have proposed the calculation of polygenic scores using a penalized regression approach using summary statistics and examined its performance in simulation experiments. Our proposed approach, 

~~~
lassosum
~~~

, in general appeared to give better prediction than *p*-value thresholding with or without clumping as well as the recently proposed LDpred, for which we failed to demonstrate the claimed superior performance over *p*-value thresholding. Clumping was beneficial for *p*-value thresholding in most scenarios but not for 

~~~
lassosum
~~~

. In some scenarios, clumping actually decreases the predictive power of *p*-value thresholding, such as in our simulations with *p*(causal) = 0.1 and *n* = 10, 000.

Compared with LDpred, we showed that 

~~~
lassosum
~~~

 is not only more accurate but also a lot faster. Running 

~~~
lassosum
~~~

 on a reference panel of around 300,000 SNPs and 1,000 individuals typically takes only several minutes without parallel processing. Even when using a reference panel with 8 million SNPs and 500 participants, 

~~~
lassosum
~~~

 took around 15 minutes without parallel processing for each value of *s*. The time taken was similar to that for clumping in PLINK 1.9 and therefore 

~~~
lassosum
~~~

 had similar speed to clumping and *p*-value thresholding when run with a small reference sample size. Increasing the sample size of the reference panel will generally increase prediction accuracy also, although this comes at a cost of exponentially increasing running times. In our simulations we found that gains in prediction accuracy from a larger reference panel were usually modest. We are currently working on a parallel implementation of 

~~~
lassosum
~~~

 and this should be available by the time the article is accepted for publication.

**Figure 2:**
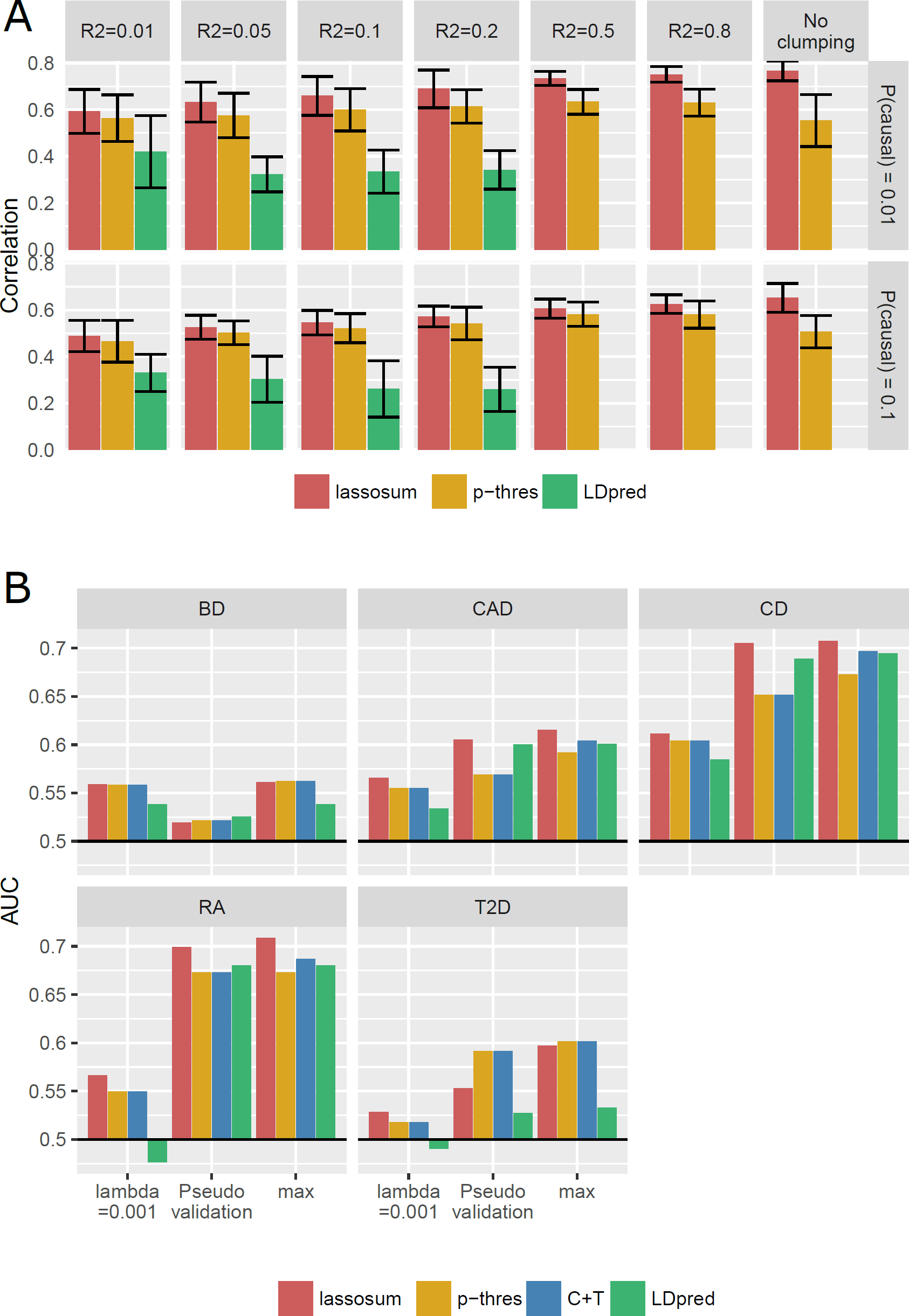
(A) Performance of 

~~~
lassosum
~~~

 in a large simulated dataset with *n* = 200, 000 using different clumping levels in relation to *p*-value thresholding and LDpred. Mean and standard deviation of the AUC of the PGS with the true disease status are plotted. (B) Performance of 

~~~
lassosum
~~~

 vs. other methods when using real summary statistics data from meta-analyses. Predictive accuracy was assessed by prediction in the WTCCC dataset after the contribution from WTCCC was removed from the summary statistics. p-thres: *p*-value thresholding without clumping, C+T: *p*-value thresholding with clumping.

Another contribution from this paper is the method of pseudovalidation, which can be applied to any PGS method that requires a tuning parameter. We showed that it is effective in selecting a parameter value that is close to the optimum. Not surprisingly, having a validation dataset with phenotype data generally provides an even more reliable method for selecting the tuning parameter. However, in the event where this is unavailable, pseudovalidation offers an alternative. Recently, polygenic scores were often used to assess genetic correlation between two diseases. Often times, the tuning parameter (or *p*-value threshold) used in the polygenic scores was chosen by maximising over the correlation of the PGS with another disease (e.g. Krapohl *et al*., 2015). We have not examined the performance of using this approach to select the tuning parameter, although it is likely that there will be bias in estimation of correlations due to winner’s curse.

Although we have focused on the performance of 

~~~
lassosum
~~~

 as a method, we note that it is more generally an instance of penalized regression. Potentially other penalties can be used in place of 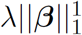 in equation (2) that can lead to even better prediction. We chose the LASSO penalty because of its simplicity. Other similar methods that can also be solved using the fast coordinate descent method of Friedman *et al*. (2007) include the non-negative garotte, LAD-LASSO, and Grouped LASSO.

Some limitations of the present study are worth bearing in mind when considering these results. For example, summary statistics may be inflated due to population stratification in the data where they are generated. As summary statistics are often derived from meta-analyses, it is also possible that there is underlying heterogeneity in effect sizes. How these impact PGS calculation is currently unknown.

Recently, methods for conducting GWAS have moved beyond the single-disease paradigm. Often, multiple related diseases are analysed together to give improved power for detection of GWAS signals (Korte *et al*., 2012; Andreassen *et al*., 2013; Zhou and Stephens, 2014; Chung *et al*., 2014; Li *et al*., 2014). Frequently, these new methods operate in the Bayesian framework resulting in Bayes factor or posterior probability of associations (or alternatively local false discovery rates) for each SNPs. In principle, we can translate these into *p*-values (Stephens and Balding, 2009) and thus make use of additional information to improve PGS predictive performance. Likewise additional information gained in the consideration of functional annotations of the genome (Schork *et al*., 2013; Pickrell, 2014; Kichaev *et al*., 2014) can be incorporated similarly. The simplicity of 

~~~
lassosum
~~~

 makes it an ideal framework from which more complex methods can be developed.

## Acknowledgments

We would like to thank Dr. Johnny S. H. Kwan for pointing out to us the work by Strimmer (2008). We would also like to thank two annoymous referees for their comments in improving this paper. We acknowledge financial support from the Hong Kong Research Grants Council General Research Fund [776513M, HKU 776412M, 17128515], the Hong Kong Research Grants Council Theme-Based Research Scheme [T12-705/11, T12/708/12N, T12C-714/14-R], the National Science Foundation of China - Research Grants Council of Hong Kong [N HKU736/14], and the European Community Seventh Framework Programme Grant on European Network of National Schizophrenia Networks Studying Gene-Environment Interactions (EU-GEI).

